# Functional phenotyping of genomic variants using multiomic scDNA-scRNA-seq

**DOI:** 10.1101/2024.05.31.596895

**Authors:** Dominik Lindenhofer, Julia R. Bauman, John A. Hawkins, Donnacha Fitzgerald, Umut Yildiz, Jan M. Marttinen, Moritz Kueblbeck, Judith B. Zaugg, Kyung-Min Noh, Sascha Dietrich, Wolfgang Huber, Oliver Stegle, Lars M. Steinmetz

## Abstract

Genomic variation ranging from single nucleotide polymorphisms to structural variants can impact gene function and expression, contributing to disease mechanisms such as cancer progression. The systematic study of this variation is hindered by inefficient precision editing tools making it challenging to confidently link genotype and gene expression in pooled screens. Additionally, assessing heterogenous variants in primary tumor samples at scale is difficult with current single-cell technologies. We developed droplet-based multiomic targeted scDNA-scRNAseq (SDR-seq) to precisely link genotypes with gene expression profiles in high-throughput. SDR-seq simultaneously assesses up to 480 RNA and gDNA targets with high coverage and sensitivity across thousands of cells. Using SDR-seq, we associate coding and non-coding variants with distinct gene expression profiles in human iPSCs. Furthermore, we demonstrate that in primary B-cell lymphoma samples, cells with a higher mutational burden exhibit elevated B-cell receptor signaling and tumorigenic gene expression. SDR-seq has broad potential for gaining functional insights into regulatory mechanisms encoded by genetic variants at diverse loci, advancing our ability to study gene expression regulation and its implications for disease.

## Introduction

Genomic variation in both coding and non-coding regions of the genome drives human population differences and disease in both mono-genetic and complex manners^1–3^. Genetic loss-of-function screening of coding genes and CRISPRi/CRISPRa screens in non-coding regions have provided valuable insights into disease mechanisms. However, they do not take into account precise genomic variation potentially masking more complex cellular disease phenotypes that are caused by individual variants^4–7^. Existing precision genome editing tools to introduce variants have limited efficiency and variable editing outcome in mammalian cells^8–10^. This makes it difficult to use gRNAs as a proxy to annotate the variant perturbation in pooled screens. Exogenous introduction of sequence variants, via episomal massively parallel reporter assays for non-coding variants or ORF expression for coding sequences, allows for high-throughput screening of sequences and their variants for functional effects, but lacks endogenous genomic position and sequence context^11–14^. These limitations hinder a systematic study of endogenous genetic variation and its impact on disease-relevant gene expression.

To confidently link precise genotypes to gene expression in its endogenous context, a combined single-cell RNA and gDNA assay is required to directly assess variants and gene expression in the same cell. Current technologies for simultaneous high-sensitivity readout of both RNA and gDNA are well-based and laborious with limited throughput^15–18^. High-throughput droplet-based or split-pooling approaches can measure thousands of cells simultaneously but lack combined high-sensitivity and tagmentation-independent readout of RNA and gDNA^19–21^. Here we developed targeted droplet-based scDNA-scRNA-seq (SDR-seq), a scalable and sensitive method to screen genetic variation in high-throughput, linking it to gene expression and distinct cellular states.

## Results

### Droplet-based scDNA-scRNA-seq (SDR-seq)

We developed SDR-seq to simultaneously measure RNA and gDNA targets in the same cell with high coverage across all cells. The assay combines an *in-situ* reverse transcription (RT) step of fixed cells with a multiplexed PCR in droplets based on the Tapestri technology from MissionBio (Fig. 1a). Cells are dissociated into a single-cell suspension, fixed, permeabilized and subjected to an *in-situ* RT using custom polydT primers. This step adds a unique molecular identifier (UMI), a sample barcode (sample BC) and a capture sequence (CS) to cDNA molecules. Cells containing cDNA and gDNA are loaded on the Tapestri machine where cells are lysed after a first droplet generation, treated with proteinase K and mixed with reverse primers for each intended gDNA or RNA target. This is followed by a second droplet generation where the first droplet is fused into a droplet containing PCR reagents, forward primers containing a CS overhang and a cell barcoding bead containing oligos with a distinct cell barcode per bead and a matching CS. A multiplexed PCR amplifies both gDNA and RNA targets in each droplet. Cell barcoding occurs by complementary CS overhangs of the PCR amplicons and oligos from the cell barcoding bead. Emulsions are broken after the PCR and sequencing ready libraries generated. Distinct overhangs on reverse primers containing either R2N (gDNA – Nextera R2) or R2 (RNA – TruSeq R2) enable to generate separate NGS libraries for gDNA and RNA. This enables to sequence these separate NGS libraries at their required specification and sequencing depth: (i) full-length to entirely cover variant information on gDNA targets, (ii) transcript and barcode information (cell BC, sample BC and UMI) for RNA targets.

**Fig. 1.**
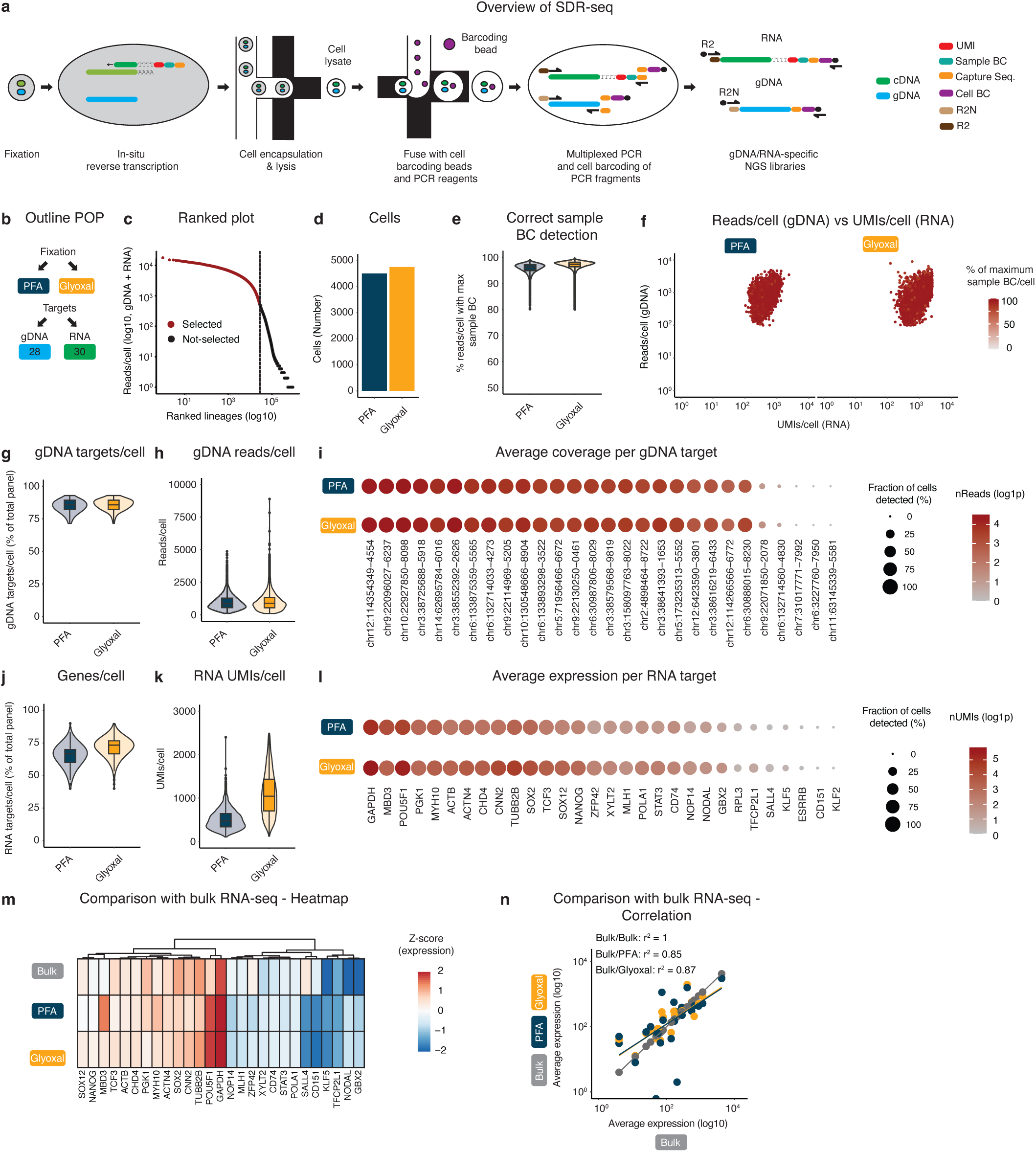
SDR-seq links gDNA variants and gene expression in single cells. **a,** Overview of targeted scDNA-scRNA-seq (SDR-seq) method. Main steps involve fixation of a single-cell suspension and *in-situ* reverse transcription (RT) followed by a multiplexed PCR within individual droplets. Both RNA and gDNA targets are amplified at the same time in each droplet. R2N (Nextera) or R2 (TruSeq) overhangs on reverse primers enable separate NGS library generation for gDNA and RNA. UMI, unique molecular identifier; BC, barcode. **b,** Outline of proof-of-principle (POP) experiment. Fixation conditions and number of gDNA/RNA targets are indicated. **c,** Knee plot of ranked cells by sequencing depth. **d,** Number of cells found per fixation condition. **e,** Correct sample BC detection per cell. Read count for maximum sample BC found was divided by the total amount of RNA reads/cell. **f,** gDNA reads/cell vs RNA UMIs/cell per fixation condition. Color indicates % reads/cell with max sample BC. **g, h,** Number of gDNA targets/cell (g) and gDNA reads/cell (h). **i,** Individual gDNA targets are shown per fixation condition. Size indicates fraction of cells detected in. Color indicates read coverage. **j, k,** Number of genes per cell (j) and RNA UMIs/cell (k). **l,** Individual gDNA targets are shown per fixation condition. Size indicates fraction of cells detected in. Color indicates UMI coverage. **m,** Comparison of expressed genes to bulk RNA-seq data. Z-score, data is scaled by row. **n,** Pearson correlation of expressed genes to bulk RNA-seq data.

To test SDR-seq we performed a proof-of-principle (POP) experiment aiming to amplify a small number of both gDNA (28) and RNA (30) targets in human induced pluripotent stem cells (iPSCs) (Fig. 1b). As fixation is a critical step to perform *in-situ* RT, we tested two different fixation conditions – PFA and glyoxal. PFA is commonly used in *in-situ* RT reactions but can have detrimental effects on gDNA and RNA quality as it crosslinks nucleic acids^22^. Glyoxal does not crosslink nucleic acids and is therefore expected to yield a more sensitive readout^23,24^. For simplicity, overhangs for reverse primers for gDNA and RNA were the same (R2N) in this POP experiment (Extended Data Fig. 1a). After SDR-seq and filtering for high-quality cells, we obtained around 9,500 cells from a single Tapestri run (Fig. 1c, d, Extended Data Fig. 1b-e). Cells were equally distributed over the two fixation conditions, and the fixation condition of each cell could be discriminated using distinct sample BCs during *in-situ* RT. On average over 95 % of reads/cell mapped to the correct sample BC (Fig. 1e, f). For downstream analysis contaminating reads were removed per cell.

gDNA target coverage is expected to be uniform across cells as each cell has the same input of gDNA. 23 out of 28 gDNA targets (82 %) had high coverage and were found in the vast majority of cells (Fig. 1g-i). Little difference was observed in gDNA target detection and coverage using either PFA or glyoxal (Extended Data. Fig. 1f, h). RNA target coverage is expected to be varying as targets were chosen over a range of expression values and cells exhibit transcriptional variability. Indeed, individual RNA targets showed varying levels of expression while some were expressed only in a fraction of the cells (Fig. 1j-l). The number of RNA targets detected and UMI coverage increased when using glyoxal as compared to PFA for fixation (Extended Data Fig. 1g, i). Ubiquitously expressed housekeeping or iPSC-maintenance genes were detected in all cells, while other genes showed specific expression only in a subset of cells, in line with published data (Extended Data Fig. 2)^25^. Comparing bulk RNA-seq data of human stem cells to pseudo bulked SDR-seq gene expression showed comparable levels of expression for the vast majority of targets with high correlation (Fig. 1m, n). These results demonstrate that SDR-seq is capable of a highly sensitive readout of many DNA and RNA targets in thousands of single cells in a single experiment with the potential to link those modalities in a high-throughput fashion.

### SDR-seq is scalable to detect hundreds of targets with high confidence

Next, we tested if SDR-seq is scalable to detect hundreds of gDNA and RNA targets simultaneously. We designed an experiment with panel sizes of 120, 240 and 480 targets consisting of equal parts of gDNA and RNA targets in iPSCs (Fig. 2a). To allow for cross comparison between panels, 60 gDNA and 30 RNA targets were shared between panels. To adjust for differences in sequencing depth, reads were subsampled for gDNA and RNA according to panel size to achieve on average an equal read coverage/cell for the shared targets between panels (Extended Data Fig. 3c-f). A gDNA target was counted as detected per cell if it was covered with at least 5 reads per cell, while an RNA target was counted if it was detected by 1 UMI per cell. We confirmed that separately prepared NGS libraries for both gDNA and RNA in this experiment mapped with high specificity to their respective references (Extended Data Fig. 3a, b). Overall, 80 % of all gDNA targets were detected with high confidence in more than 80 % of cells across all panels with only a minor decrease in detection for bigger panel sizes (Extended Data Fig. 4a-c). Detection and coverage of shared gDNA targets between panels was well correlated indicating that detection of gDNA targets is largely independent of panel size (Figure 2b, c). The minor decrease in detection rate for the bigger panel sizes predominantly affected targets with lower coverage across panels (Extended Data Fig. 4d, e, Extended Data Fig. 5a). Checking all RNA targets we observed that there was as a minor decrease in targets detected in larger panels compared to the 120 panel (Extended Data Fig. 4f-h). Detection and gene expression of shared RNA targets were highly correlated between all panels (Fig. 2d, e, Extended Data Fig. 4i, j) showing a robust and sensitive detection of gene expression readouts independent of panel size. Variability was predominantly observed for lowly expressed genes (Extended Data Fig. 5b).

**Fig. 2.**
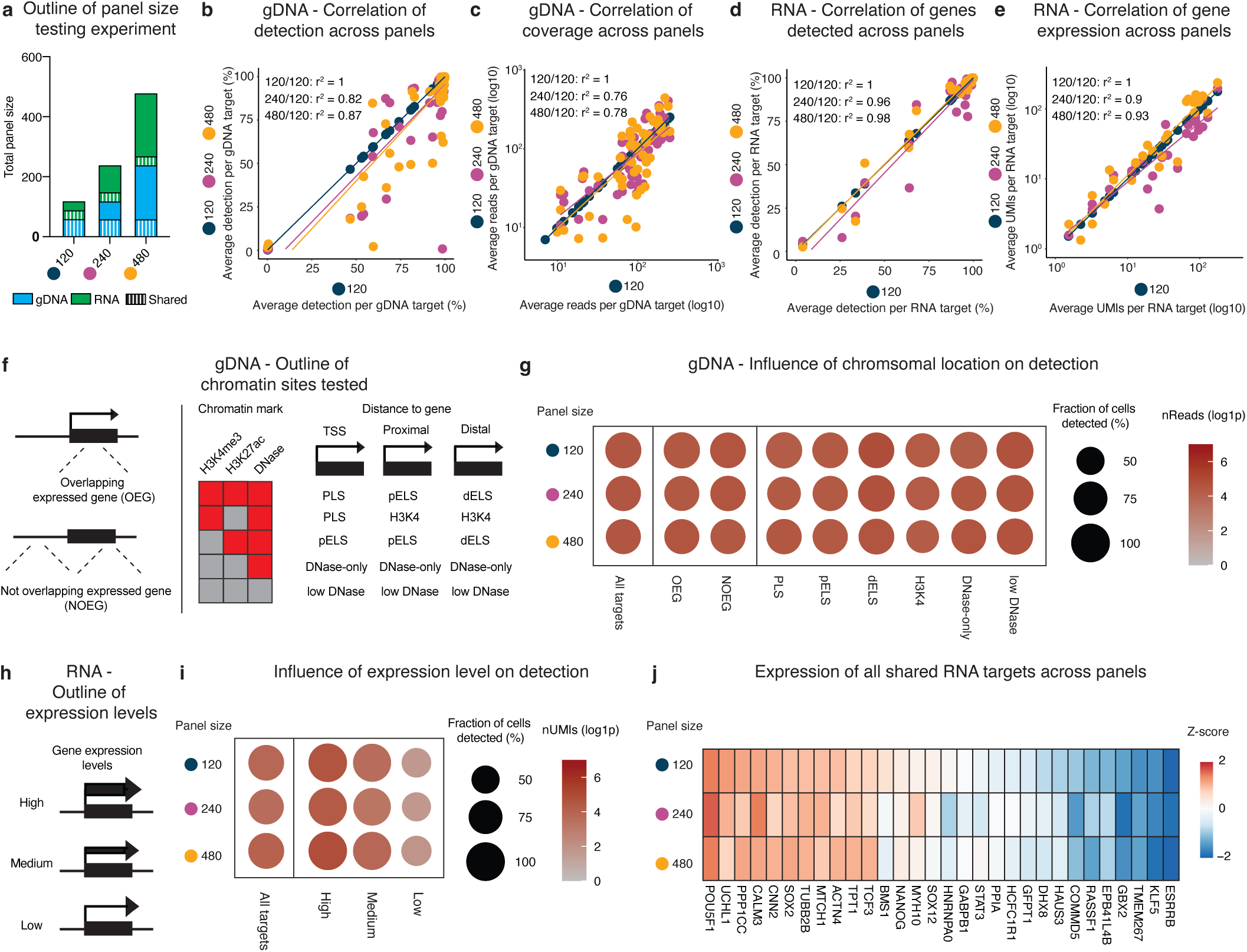
SDR-seq scales to hundreds of targets simultaneously. **a,** Outline of panel size testing experiments. gDNA and RNA targets are equal within panels; Shared targets are indicated. **b, c,** Pearson correlation of detection (b) and coverage (c) of shared gDNA targets between panels. **d, e,** Pearson correlation of detection (b) and coverage (c) of shared genes between panels. **f,** Outline of chromatin sites tested. OEG, overlapping expressed gene. NOEG, not overlapping expressed gene. Combination of chromatin marks detected and the relative distance to a gene define regulatory elements. PLS, promotor-like sequence. pELS, proximal enhancer-like sequence. dELS, distal enhancer-like sequence. **g,** Detection of different chromatin sites between panels. Size indicates fraction of cells detected in. Color indicates read coverage. **h,** Outline of expression levels tested. **i,** Detection of genes with different expression levels between panels. Size indicates fraction of cells detected in. Color indicates read coverage. **j,** Heatmap of expression of all shared genes between panels. Z-score, data is scaled by row.

To check whether chromosomal context has an influence on gDNA detection using SDR-seq, we included targets sites among the shared panels that were either overlapping (OEG) or not overlapping expressed genes (NOEG). Additionally, we tested for different chromatin marks and states (H3K3me3, H3K27ac, DNase sensitive) indicating different genomic regulatory elements depending on their proximity to the TSS (Fig 2f)^26^. We did not observe a strong impact on detection and coverage across panels depending on OEG or NOEG location (Fig. 2g). Importantly there was no regulatory element that showed a systematic bias to be detected, while also sites with low-DNase signal were confidently recovered.

Genes were chosen based on a range of expression dividing them into high, medium and lowly expressed genes (Fig. 2h). While highly and medium expressed genes were detected in almost all cells, lowly expressed genes showed a decreased detection rate across panel sizes (Fig. 2i). This is in line with published data, where some genes are not found to be expressed in all cells in iPSCs^25^. Overall expression levels of shared genes were highly similar across the different panel sizes tested (Fig. 2j).

SDR-seq is thus scalable to assay hundreds of gDNA and RNA targets simultaneously with high reproducibility and sensitivity across different panel sizes independently of chromatin state and expression level. This makes it a versatile tool to both assay variants in hundreds of loci in single cells, while linked gene expression changes and cell identity markers can be measured.

### SDR-seq confidently detects gene expression changes

Genomic variants can increase or decrease gene expression, but effect sizes are often expected to be small. Therefore, a sensitive readout of these gene expression changes is critical. To test if SDR-seq sensitively measures gene expression changes, we designed a CRISPRi experiment consisting of non-targeting control gRNAs (NTC), gRNAs targeting high-confidence expression quantitative trait loci (eQTLs), gRNAs targeting the transcription start site (TSS) of genes predicted to be affected by those eQTLs (CRISPRi controls) and gRNAs that target the gene body of a transcript where a possible STOP codon could be introduced via prime editing (STOP controls) (Fig. 3a). Human iPSCs expressing the CRISPRi transgene from the safe-harbor locus AAVS1 were infected with a lentiviral CROP-seq gRNA library, selected via FACS and SDR-seq was performed (Fig. 3b). The SDR-seq primer panel amplified the gDNA sites of the eQTLs together with associated transcripts, the viral CROP-seq transcript to assign cells to gRNAs and multiple housekeeping genes to normalize data independent of perturbation effects. Cells were successfully assigned to gRNAs (75 %) resulting in an average coverage of 30 cells/gRNA (Extended Data Fig. 6a, b). NTC gRNAs did not show a significant effect on any of the genes measured, while the vast majority (95 %) of CRISPRi control gRNAs targeting the TSS of a gene showed a strong effect on the expression levels of the associated target gene (Fig. 3c). Seven eQTL gRNAs (24 %) and three STOP control gRNAs (60 %) showed a significant effect on target gene expression. Significantly scoring eQTL and STOP control gRNAs were located within a two kb window of the TSS indicating a direct inhibitory effect of CRISPRi comparable to the CRISPRi control gRNAs (Extended Data Fig. 6c). This shows that SDR-seq can confidently detect gene expression changes mediated by CRISPRi targeting the TSS. Additionally, this data highlights the importance of directly assessing variants in proximity to TSS to check their impact on gene expression rather than approximating such effects with CRISPRi.

**Fig. 3.**
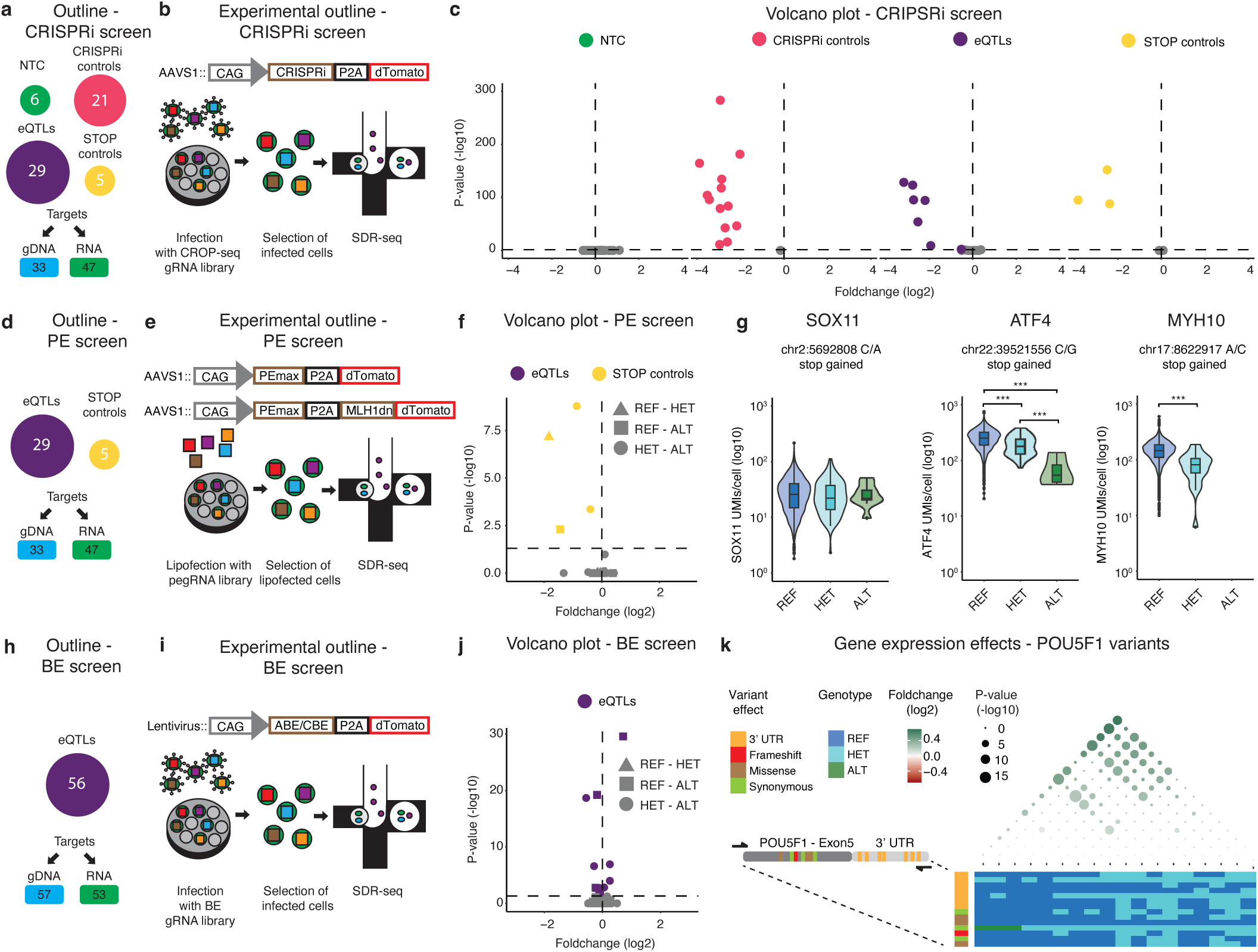
SDR-seq is sensitive to detect gene expression changes and link them to variants. **a,** Outline of the CRISPRi screen. NTC, non-targeting control. eQTL, expression quantitative trait loci. **b,** Experimental outline of the CRISPRi screen. **c,** Volcano plot for CRISPRi screen with different gRNA classes indicating foldchange and *P*-value. Significant hits (*P*-value < 0.05) are colored. For NTCs all genes measured are shown. For other gRNA classes only the intended target for each gRNA is shown. **d,** Outline of prime editing (PE) screen. **e,** Experimental outline of the PE screen. **f,** Volcano plot for PE screen with different gRNA classes indicating foldchange and *P*-value. Significant hits (*P*-value < 0.05) are colored. Comparison between the different alleles is shown as shapes. **g,** Alleles and gene expression for SOX11, ATF4 and MYH10 STOP controls are shown. \*\*\**P* < 10^−4^ by MAST. **h,** Outline of base editing (BE) screen. **i,** Experimental outline of the BE screen. **j,** Volcano plot for different gRNA classes indicating foldchange and *P*-value. Significant hits (*P*-value < 0.05) are colored. Comparison between the different alleles is shown as shapes. **k,** Variants in POU5F1 locus and their impact on gene expression is shown. Impact of variant is color coded. REF, HET and ALT alleles are shown for each genotype. Foldchange between combination of variants is indicated in color (green), *P*-value as size (-log10).

Next, we aimed to directly install eQTL variants and measure their effect sizes on gene expression. We generated two iPSCs cell lines that express a transgene to enable prime editing (PE), with or without the simultaneous expression of a dominant negative regulator of the mismatch repair pathway that was shown to increase editing efficiency (PEmax or PEmax-MLH1dn)^10^. We tested these PE iPSCs with a fluorescent lentiviral reporter system that enables measurement of editing efficiencies via the reconstitution of a non-functional EGFP (Extended Data Fig. 6d). We observed around 50 % editing efficiency after lipofection of a prime editing gRNA (pegRNA) that repairs the EGFP fluorescent protein (Extended Data Fig. 6e) indicating the system has the potential to enable editing in iPSCs. Next, we lipofected these validated PE iPSCs with a pegRNA library to introduce the same eQTLs as in the CRISPRi screen, or STOP codons to assay nonsense mediated decay. We enriched cells that were lipofected with flow cytometry and performed SDR-seq (Fig. 3e). We observed limited editing efficiency with both PE cell lines confounding the interpretation of many variants (Extended Data Fig. 7a, b). Variants could be confidently called discriminating between reference (REF), heterozygous (HET) and alternative (ALT) alleles (Extended Data Fig. 7c-e). We only observed significant gene expression changes for the STOP controls (Fig. 3f). Depending on the position of the STOP codon within the transcript, effects of nonsense mediated decay on transcript levels can vary^27^. For SOX11 we observed no changes on transcript levels, while STOP codons introduced in ATF4 and MYH10 had a significant impact on gene expression (Fig. 3g).

In addition to installing eQTLs with PE, we tested the use of base editors (BE) in human iPSCs. We selected 56 high confidence eQTLs to be installed with either ABE8e or CBE base editors (Fig. 3h). After introducing gRNA libraries into iPSCs, cells were selected and SDR-seq performed (Fig. 3i). We found several eQTL variants with a significant effect on target gene expression (Fig. 3j). Additionally, we measured the effect of non-BE associated mutations using SDR-seq. Human iPSCs accumulate somatic mutations during cell culturing, while they undergo constant competitive selection for variants that are advantageous in culture conditions^28^. We found a synonymous variant in the 3’ end of POU5F1, a critical factor for iPSCs maintenance, that showed a significant effect on gene expression (Extended Data Fig. 8a). However, after assessing variants that may have accumulated during culturing along the entire amplicon of POU5F1, we found that certain combinations of variants showed different effects on POU5F1 expression (Figure 3k, Extended Data Fig. 8b). In particular a set of variants in the 3’ UTR was associated with significantly different transcript levels. This highlights the importance of directly assessing variants at the locus of interest to get a better resolution on their impact on gene expression.

SDR-seq can confidently detect variants on a single cell level and associate them with gene expression differences with sensitivity to detect even subtle changes. This is the case even under conditions of limited editing efficiency in our experiments, which confound the interpretation of many tested eQTLs.

### Intra-tumor variants associate with distinct tumorigenic expression patterns in B-cell lymphomas

Linking variants to gene expression profiles remains challenging in primary samples and is of special interest to study cancer pathogenesis and evolution. B-cell non-Hodgkin lymphomas are a heterogenous set of cancers of the lymphatic system which arise from distinct stages of the B-cell affinity maturation differentiation trajectory. Here, naïve B-cells (Naïve) undergoing T-cell dependent activation migrate through the dark zone (DZ) and light zone (LZ) of the germinal center where they undergo somatic hypermutation of the B-cell receptor and selection by antigen-presenting cells, followed by maturation into memory B-cells (Mem IgM or Mem IgG) and plasma cells (Plasma)^29–32^. While the cell of origin is central to the classification of B-cell lymphomas and other cancers, it was recently shown that cancer cells can retain their ability to differentiate along their innate differentiation trajectory, whereby tumors evolve multiple maturation states from the same cell of origin^33^. Meanwhile, clonal evolution posits that tumors also evolve through the accumulation of heterogenous genetic variants over time resulting in the generation of treatment resistant sub-clones^34^.

We used B-cell lymphomas to investigate how genetic variation influences gene expression and maturation states arising through differentiation within tumors. We characterized frequent variants in two follicular lymphoma (FL) primary patient samples and one germinal center subtype diffuse large B-cell lymphoma (GCB) primary patient sample using SDR-seq (Fig. 4a). Assaying between 3,600 and 8,400 cells per patient sample we could distinguish B-cells from other cells based on gene expression of the targeted RNA panel (Fig. 4b, c). B-cells of each primary patient sample formed separate clusters while cells of other identity (non-B-cells) clustered together. We leveraged a data integration approach based on mutual nearest neighbors and canonical correlation analysis to map B-cell maturation states from a published single-cell transcriptomic dataset in non-malignant reactive lymph nodes to tumor samples (Extended Data Fig. 9a-b)^33,35^. Restriction of the immunoglobulin light chain among the vast majority of B-cells in each tumor to either kappa or lambda confirmed their monoclonality and therefore malignancy (Extended Data Fig. 9c)^36^. Heterozygous (HET) or homozygous variants (ALT) that were observed in both malignant B-cells and non-B-cells indicate a somatic origin with a limited contribution to disease progression (Fig. 4d). Heterozygous or homozygous variants occurring in malignant B-cells specifically compared to non-B-cells could potentially contribute to disease progression in those cancers. The three patient samples showed a number of distinct variants while some predominately somatic variants were shared.

**Fig. 4.**
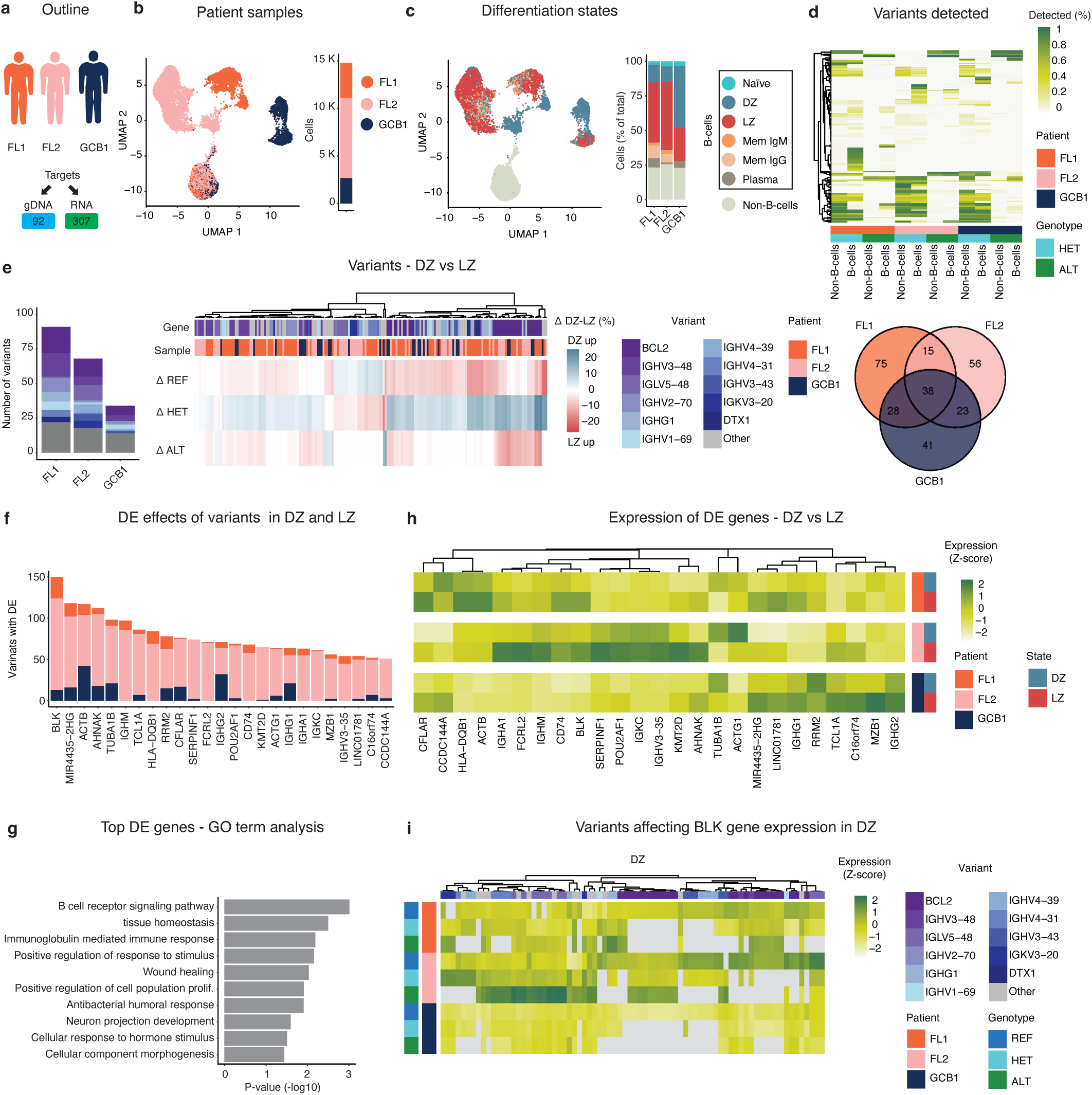
SDR-seq to profile primary non-Hodgkin lymphoma patient samples. **a,** Outline of experiment. Patient samples and target panels are indicated. **b,** UMAP highlighting the different patient samples. Number of cells for each patient sample is indicated as a bar graph. **c,** UMAP highlighting the maturation states. Number of cells within a maturation state is indicated as a bar graph as percentage of total. **d,** Variants detected in the experiment. Color indicates percentage within B-cells and non-B-cells for each variant. Patient samples and HET/ALT alleles are indicated by color. Venn-Diagram shows overlap of variants that occur with more than 5 % of frequency in each sample. **e,** Significantly differentially abundant variants comparing DZ and LZ states. Summed counts of genes which the variants map to shown in a bar graph. Δ DZ-LZ (% of the respective allele in DZ minus LZ) as percent shown in a heatmap. Genes which the variants map to and patient of origin are indicated by color. **f,** Summed counts for differentially expressed (DE) genes between variant containing and non-containing cells within both DZ and LZ for most frequent variants. **g,** GO term analysis of most frequently DE genes. **h,** Gene expression of most frequently DE gene across patient samples in LZ and DZ states. Color indicates expression (Z-score, data is scaled by column), patient samples and maturation state. **i,** BLK expression in LZ and DZ in cells with variants. Color indicates expression (Z-score, data is scaled by column), patient samples, genotype, maturation state and genes which variants map to are indicated.

Next, we focused on a comparative analysis between DZ and LZ maturation states as most cells belonged to these states (> 80 % among B-cells) and prediction scores to assign these states were highest with the targeted RNA panel used. Differential abundance testing of variants between DZ and LZ showed that most of the differentially enriched variants mapped to BCL2, an anti-apoptotic factor central to B-cell maturation in the LZ that is commonly overexpressed in B-cell lymphomas (Fig. 4e)^37^. Variants were also enriched in many immunoglobulin variable genes which are targeted during somatic hypermutation. Cells in the LZ state showed predominantly an increase in homozygous or WT alleles when compared to DZ, while the same variants showed a corresponding decrease in the DZ state compared to LZ (Fig. 4e). Next, we tested if frequent variants have an impact on gene expression in cells belonging to either LZ or DZ states. We subset cells within each state into variant containing or not-containing and performed differential gene expression (DE) testing within each state. This revealed a number of genes involved in B-cell receptor signaling and tumorigenesis frequently affected in both DZ and LZ states (Fig. 4f, g). Across patient samples, these genes showed patient-specific higher expression levels predominantly in LZ compared to DZ (Fig. 4h). The most frequently affected gene in the DE analysis was BLK, a kinase that is central to B-cell receptor signaling and maturation^38,39^. Homozygous or heterozygous variants showed elevated levels of BLK expression in the DZ state compared to WT alleles in FL patient samples, while BLK expression was either unchanged or moderately increased in the GCB patient sample (Fig. 4i). LZ cells in the geminal center (GC) can revert to the DZ upon unsuccessful binding to antigens from antigen presenting cells and thereby undergo multiple rounds of somatic hypermutations ^40^. Our data suggest that cells with a high mutational burden may have undergone more rounds of somatic hypermutation have increased B-cell receptor signaling and tumorigenic gene expression patterns to evade apoptosis induced by unsuccessful antigen binding in the LZ. This is in line with the distinct enrichment of variants in LZ compared to DZ we observe.

Using SDR-seq, we can profile variants and gene expression in a combined fashion in primary tumor samples linking cell states to mutational burden. We can assess variants that are present in malignant B-cells and non-B-cells and test for enrichment of variants in maturation states and their impact on gene expression. This reveals elevated tumorigenic and anti-apoptotic signaling in cells with higher mutational burden in distinct states within our assayed patient samples.

## Discussion

Here we developed SDR-seq to directly measure gDNA and RNA in single cells with high-throughput and sensitivity. This method uses targeted primer panels to directly assess gDNA and RNA in the same cells in droplets via a multiplexed PCR. SDR-seq achieves high coverage of gDNA targets using cost-effective sequencing, enabling precise determination of variant zygosity and linking it to sensitive gene expression readouts. This contrasts with existing split pooling or droplet-based approaches that rely on tagmentation on nucleosome depleted chromatin and require reading out the entire genome for each cell resulting in sparse data and difficulties to correctly determine zygosity of variants^19–21^. Our results demonstrate that SDR-seq can assay hundreds of gDNA and RNA targets simultaneously with high reproducibility and sensitivity across different panel sizes. The scalability and sensitivity of SDR-seq make it a versatile tool for studying a wide range of genetic variants and their effects on gene expression across diverse cell types. We can link variants in both iPSCs and primary patient samples to distinct gene expression patterns and are sensitive to detect minor gene expression changes. Advances in prime editing and pegRNA prediction tools will allow to overcome limitations we observed in this study constraining the interpretation of several eQTLs. In B-cell lymphoma patient samples, SDR-seq enabled the identification of tumor-specific variants and their associated gene expression profiles, highlighting its potential for studying intratumor heterogeneity and cancer evolution. We could associate cells with higher mutational burden to elevated B-cell receptor signaling and tumorigenic gene expression in primary B-cell lymphoma patient samples.

In future applications, SDR-seq could be combined with other readouts including a targeted protein readout or DNA methylation to provide a more holistic view of cellular regulation^41,42^. Further improvements in the gene expression readout might enable measuring larger RNA panels, or a whole transcriptome readout in parallel to a highly sensitive targeted gDNA readout for multiple loci, broadening the scope of potential applications.

SDR-seq offers a powerful, scalable, and sensitive approach to link genomic variants to gene expression in single cells and is flexible to assay both genetically engineered cell lines and primary patient tissue samples. In combination with gene editing tools it holds great potential to decipher the regulatory mechanisms that underlie endogenous variants, complementing other high-throughput approaches that assay the gene expression-to-variant link of endogenous loci or via barcoding approaches^11–14,43,44^. This method advances our ability to study gene expression regulation and its implications for disease, providing new insights that could drive the development of therapeutic strategies and enhance our understanding of complex genetic disorders.

## Acknowledgments

We thank all members of the Steinmetz lab for feedback and technical expertise. We thank the Genomics Core Facility (GeneCore) at EMBL for consultation and sequencing. We thank Jan-Philipp Mallm and the Single-cell Open Lab (scOpenLab) at DKFZ for expertise and access to the Tapestri instrument from MissionBio. We thank the Flow Cytometry Core Facility at EMBL for support and flow cytometry services. **Funding:** D.L. received funding from a HFSP postdoctoral fellowship (LT0023/2022-L). DF. received funding from a grant from the German Federal Ministry of Education and Research (SIMONA, 031L0263A).

## Author Contribution Statement

D.L. and L.M.S. designed the study, analyzed data and wrote the manuscript with input from all authors. D.L. and J.R.B. performed experiments and analyzed data with help from M.K. D.L., J.R.B., J.A.H. and D.F. performed bioinformatic analysis. U.Y. and J.M.M. provided protocols for fixation compatible with *in-situ* reverse transcription. J.B.Z., K.M.N., O.S., S.D., W.H. and L.M.S. acquired funding.

---

**Extended Data Fig. 1.**
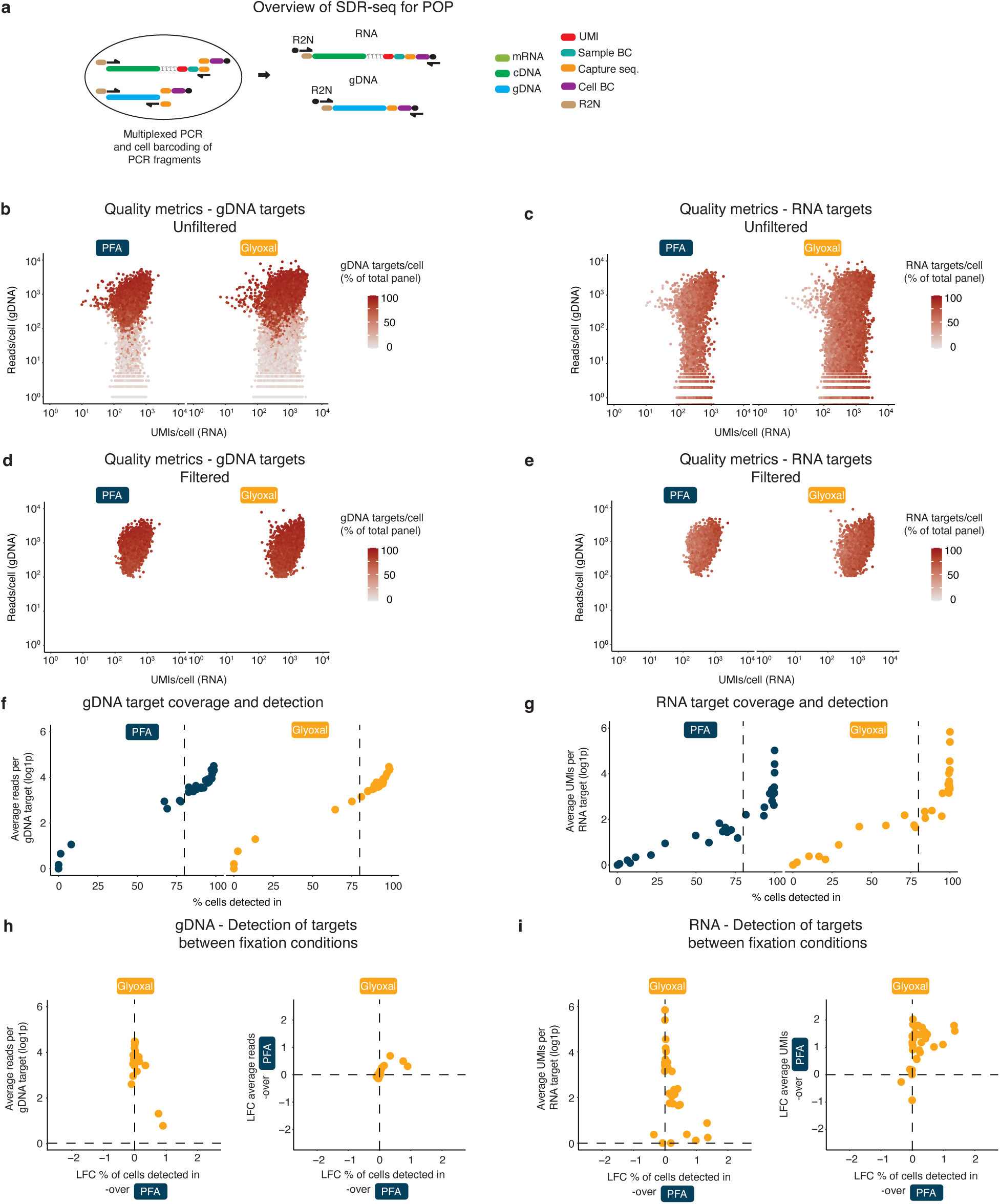
Quality controls and comparison of PFA and glyoxal fixation conditions. **a,** Overview of targeted scDNA-scRNA-seq (SDR-seq) method for proof-of-principle (POP). R2N (Nextera) reverse primer overhangs were used for both gDNA and RNA. **b-e,** Quality control plots before (b, c) and after (d, e) filtering for low quality cells. Color indicates fraction of detected gDNA or RNA targets respectively. **f, g,** Coverage and detection of each gDNA (f) or RNA (g) target. Dashed line indicates 80 % detection. **h, i,** Comparison of coverage and detection between PFA and glyoxal for each gDNA (h) and RNA (i) target. LFC, log fold change (log2).

**Extended Data Fig. 2.**
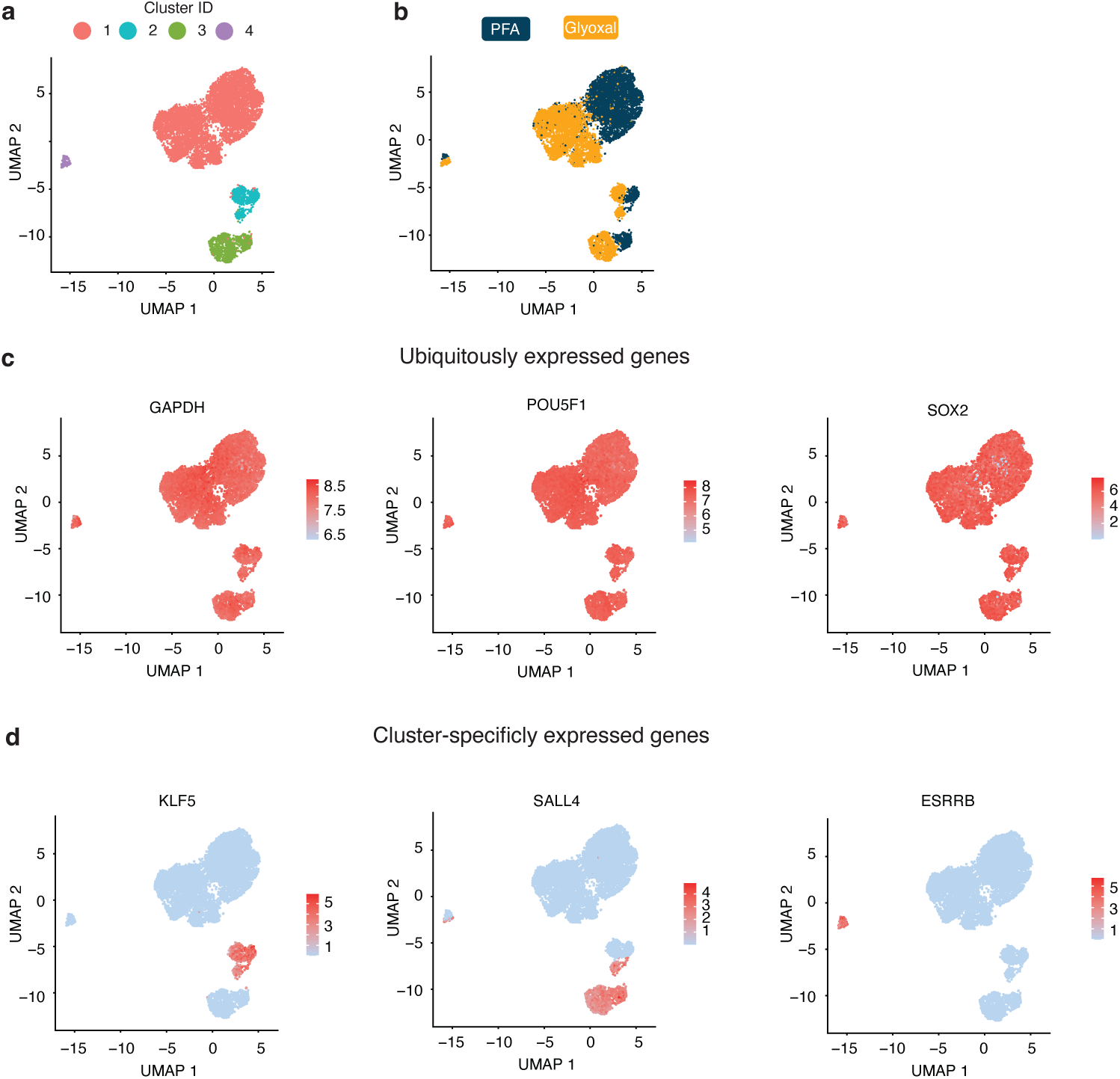
UMAPs and clustering of POP data. **a,** UMAP clustering of POP SDR-seq data. **b,** Color coding of UMAP by fixation condition. **c,** UMAP plots of ubiquitously expressed genes GAPDH, POU5F1 and SOX2. Color indicates normalized expression (log1p). **d,** UMAP plots of cluster-specifically expressed genes KLF5, SALL4 and ESRRB. Color indicates normalized expression (log1p).

**Extended Data Fig. 3.**
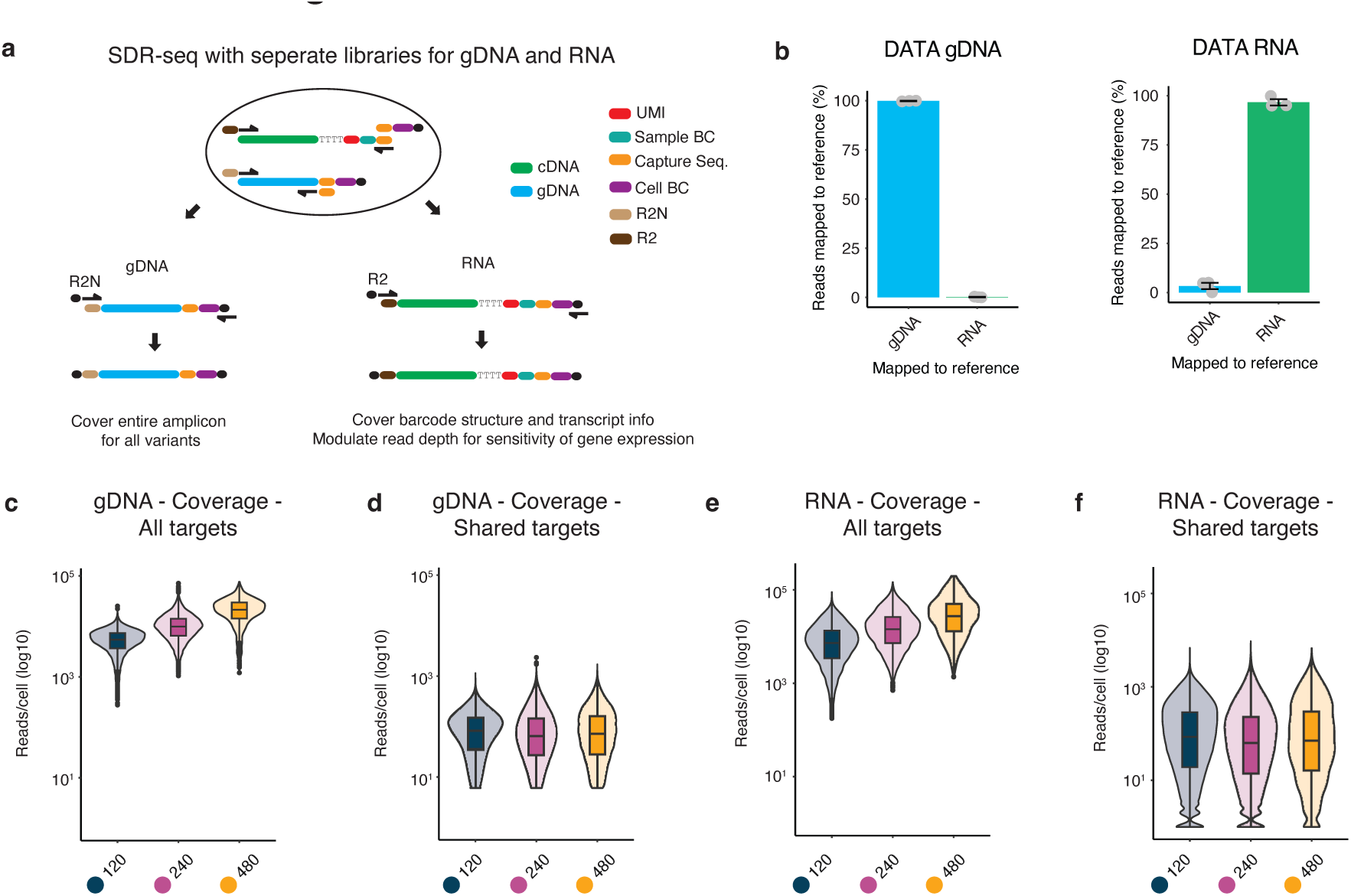
SDR-seq with separate library generation for RNA and gDNA targets. **a,** Overview of SDR-seq with separate library generation for RNA and gDNA. Modification includes a distinct R2 or R2N overhang for each library. Sequencing ready libraries can be generated using specific library primers binding to R2 or R2N, respectively. **b,** Specificity of gDNA and RNA NGS libraries. Data from gDNA or RNA libraries was mapped to either gDNA or RNA references. **c, d,** Subsampled reads/cell for all gDNA targets (c) or shared gDNA targets (d). **e, f,** Subsampled reads/cell for all RNA targets (e) or shared RNA targets (f).

**Extended Data Fig. 4.**
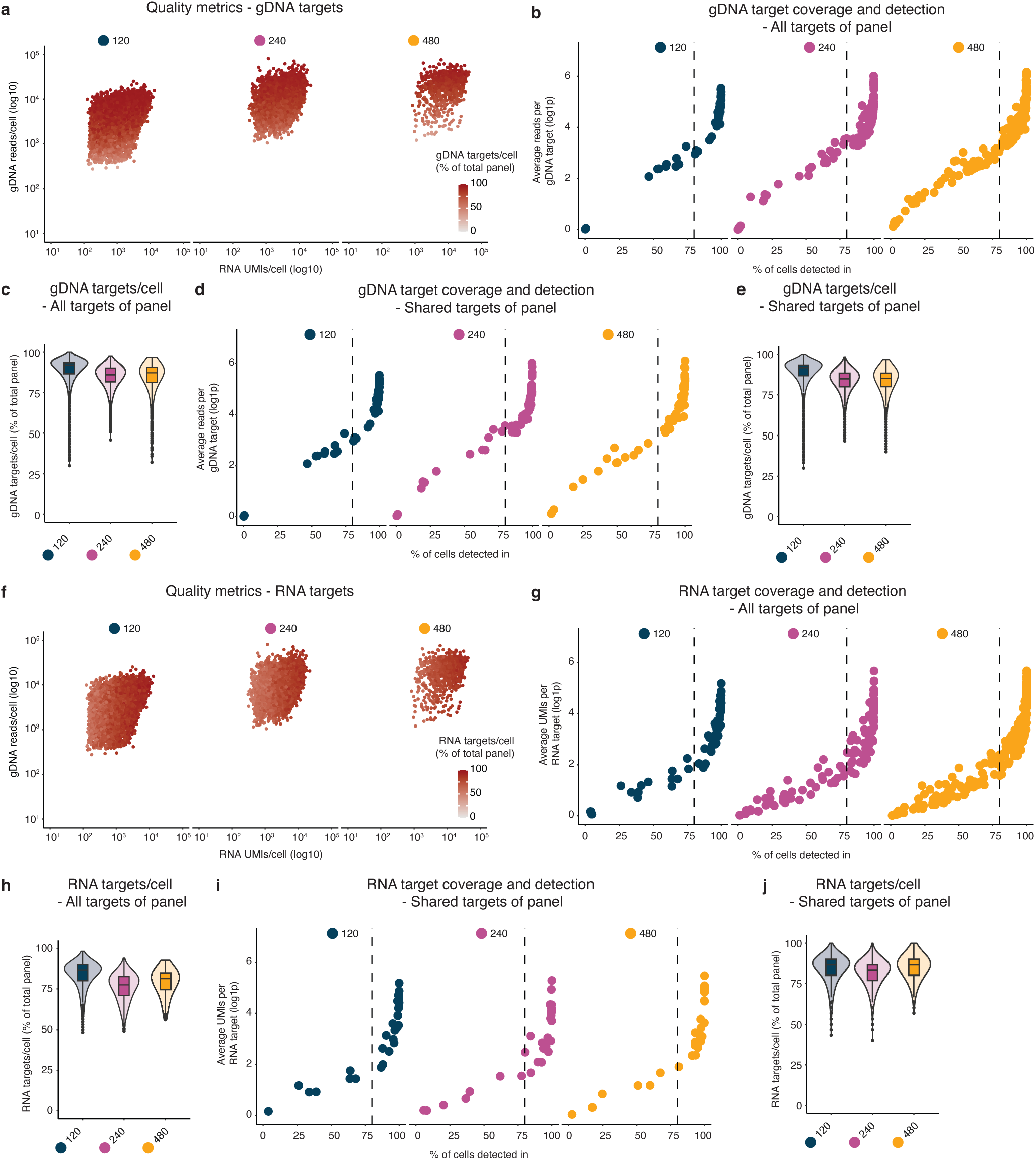
Metrics for target detection and coverage across differently sized target panels. **a,** Quality metrics of panel size experiment. Color indicates fraction of gDNA targets/cell recovered. **b,** gDNA coverage and detection for all targets across panels tested. Dashed line indicates 80 % detection. **c,** gDNA target detection per cell for all targets across panels tested. **d,** gDNA coverage and detection for shared targets across panels tested. Dashed line indicates 80 % detection. **e,** gDNA target detection per cell for shared targets across panels tested. **f,** Quality metrics of panel size experiment. Color indicates fraction of RNA targets/cell recovered. **g,** RNA coverage and detection for all targets across panels tested. Dashed line indicates 80 % detection. **h,** RNA target detection per cell for all targets across panels tested. **i,** RNA coverage and detection for shared targets across panels tested. Dashed line indicates 80 % detection. **j,** RNA target detection per cell for shared targets across panels tested.

**Extended Data Fig. 5.**
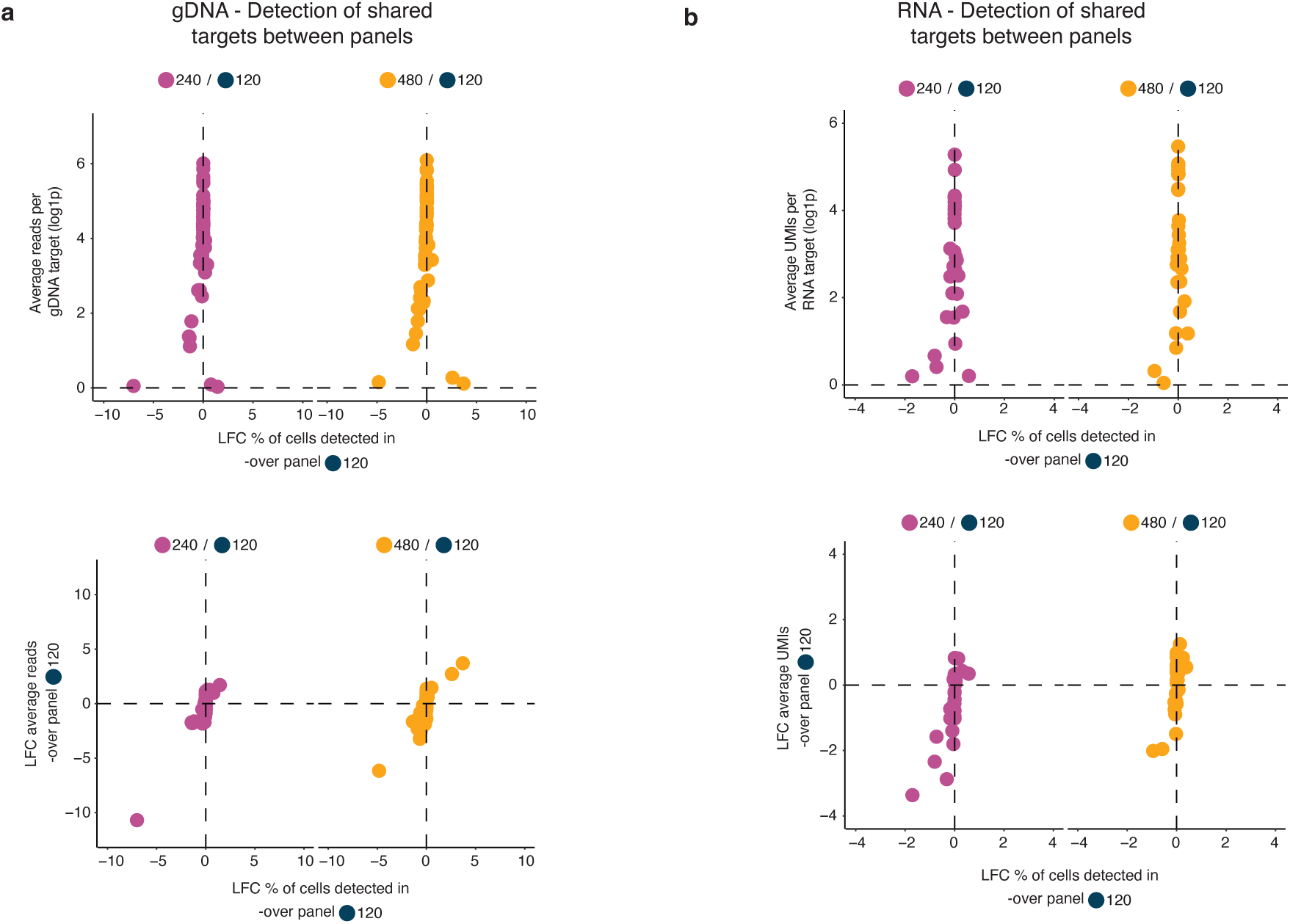
Comparison of target detection and coverage across differently sized target panels. **a, b,** Comparison of coverage and detection for shared targets across panels tested for each gDNA (a) and RNA (b) target. LFC, log fold change (log2).

**Extended Data Fig. 6.**
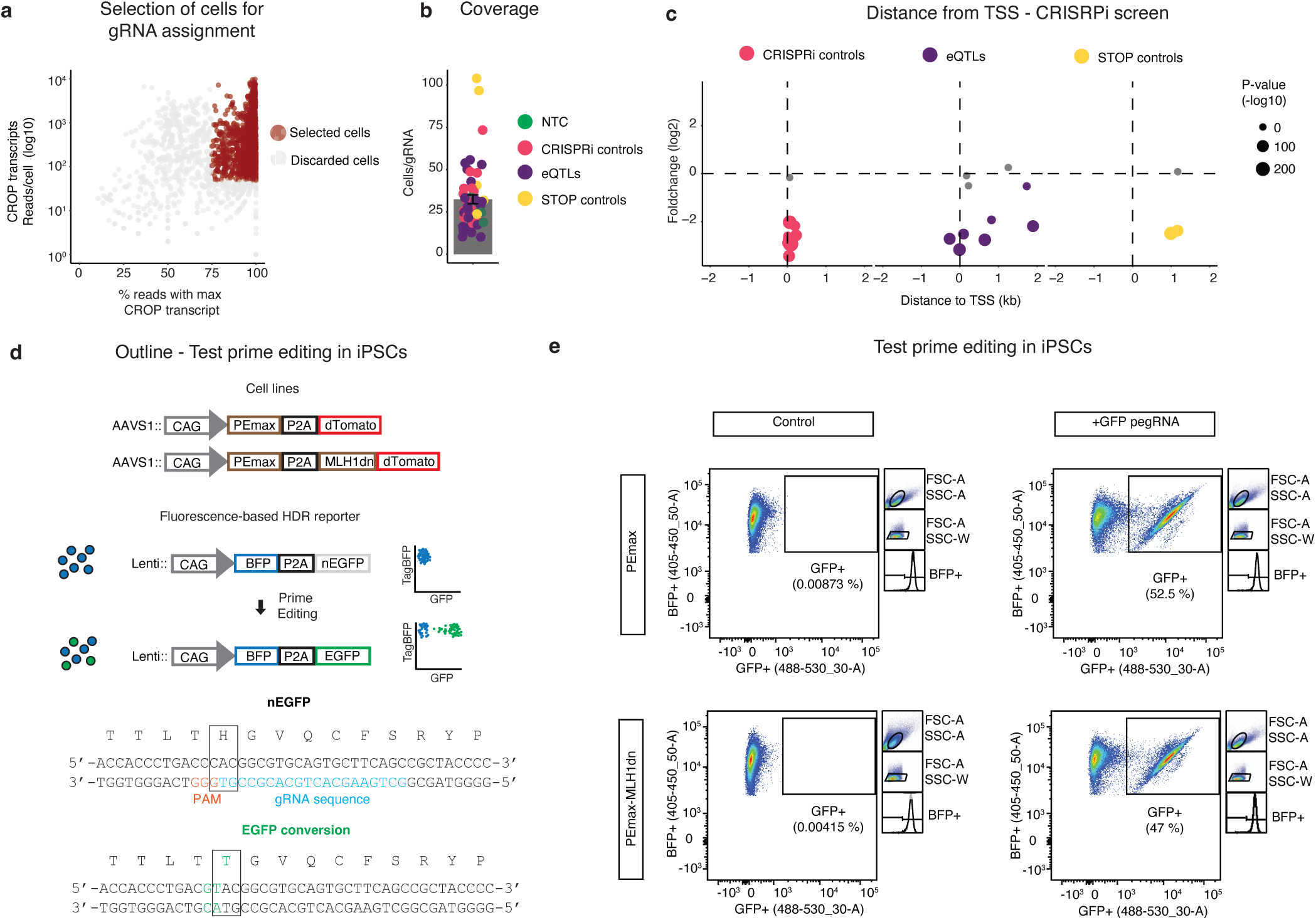
Quality metrics for CRISPRi screen and functional testing of PE iPSCs. **a,** Overview of gRNA assignment for CRISPRi screen. **b,** Coverage for each gRNA in CRISPRi screen. **c,** Relative distance of gRNA binding site to transcription start site (TSS). Positive values are after TSS (within transcript), negative values before transcript. Size indicates *P*-value. Significant hits (*P*-value < 0.05) are colored. **d,** Outline of testing for PE iPSCs. Editing can be measured by repairing a non-functional EGFP that was integrated via a lentivirus. **e,** Flow cytometry indicating editing in PE iPSCs.

**Extended Data Fig. 7.**
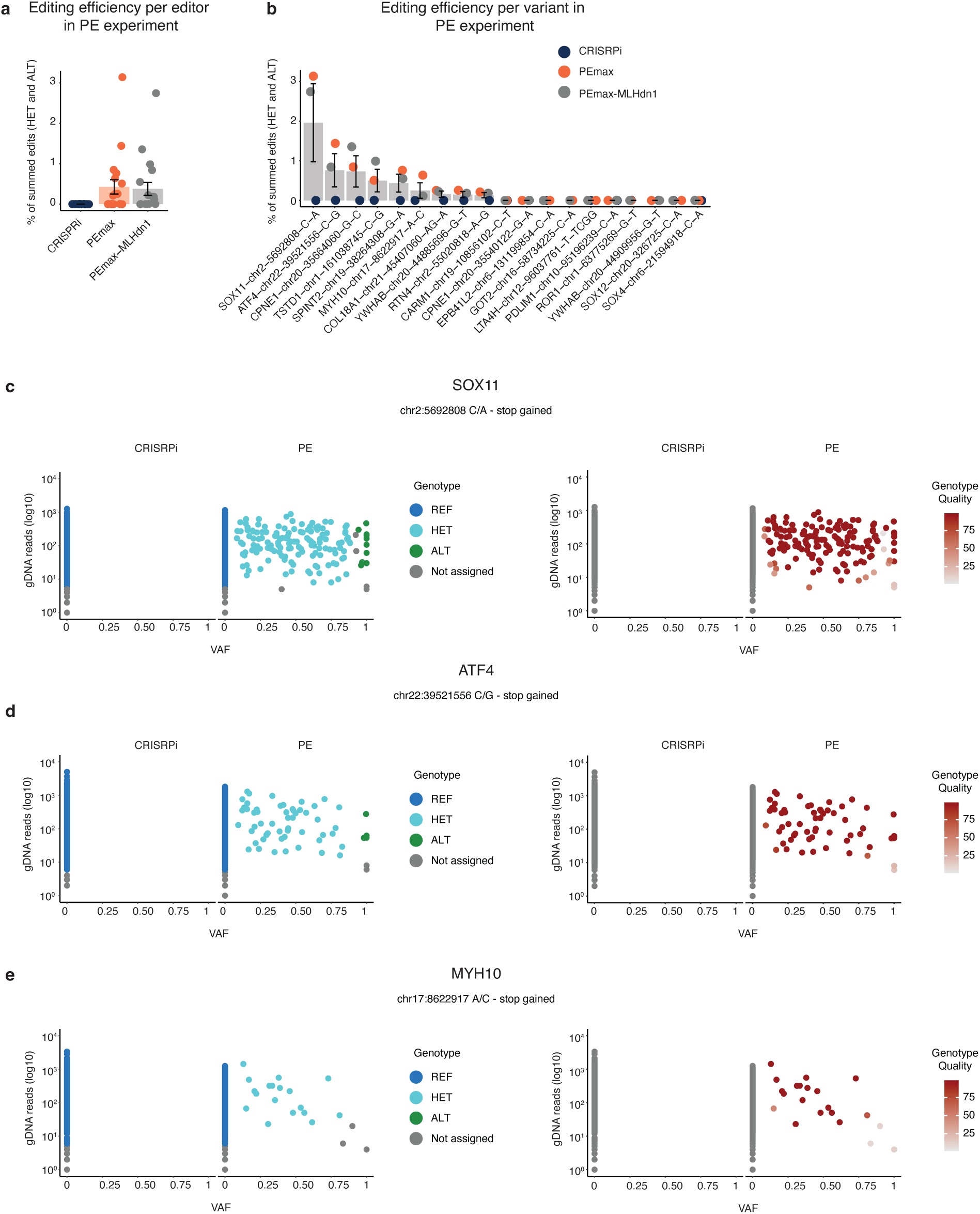
Editing efficiency and genotyping in PE screen. **a,** Editing efficiency in PE screen. CRISPRi is indicated as a control. **b,** Editing efficiency for each locus that was assessed. Loci that had somatic alleles in the PE iPCSs that were either HET or ALT are not shown. Color indicates either PE cell line or CRISPRi. **c-e,** Called genotypes and genotype quality vs. variant allele frequency (VAF) for SOX11 (c), ATF4 (d) and MYH10 (e).

**Extended Data Fig. 8.**
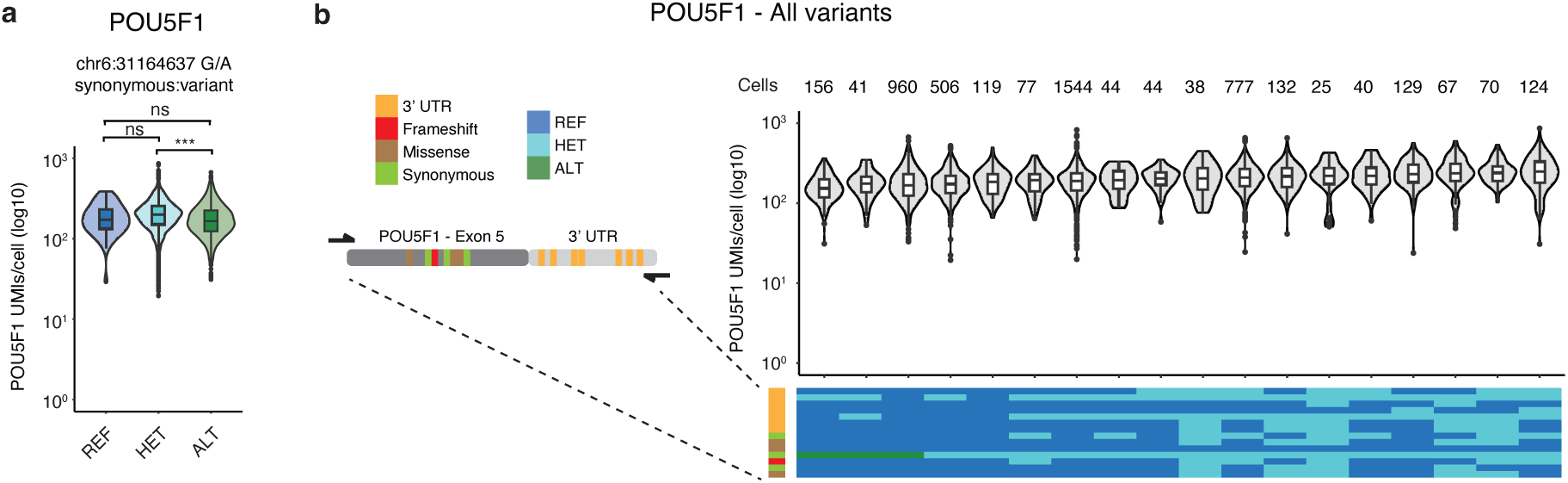
POU5F1 locus in BE screen. **a,** Intended edit to be introduced by base editing gRNA is shown at the POU5F1 locus and its impact on gene expression. **b,** All measured variants in combination along the measured gDNA site shown for POU5F1 with their impact on gene expression.

**Extended Data Fig. 9.**
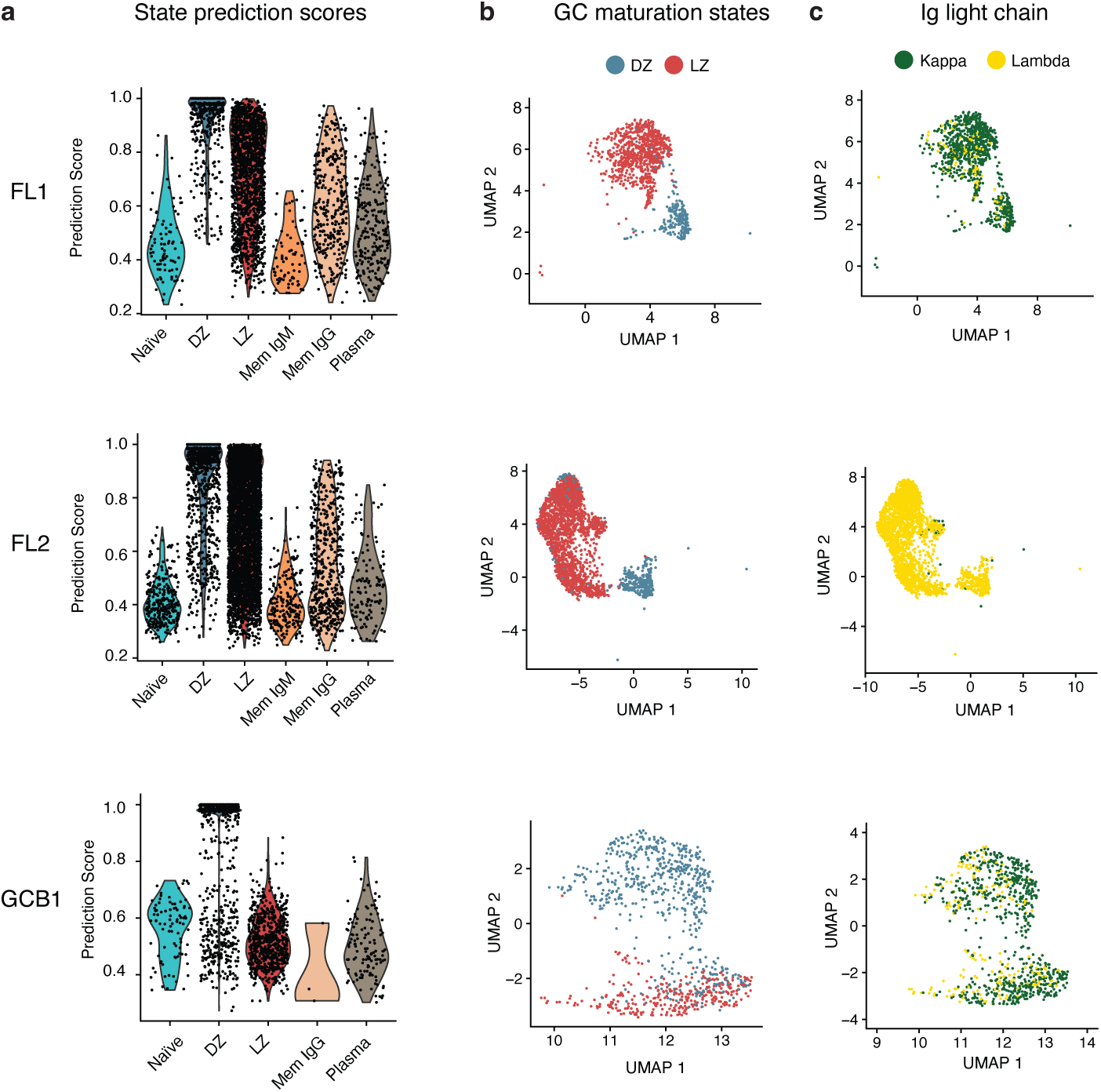
Assignment of maturation states in non-Hodgkin lymphoma patient samples. **a,** Prediction scores of cell types assignment using a reference dataset^33,35^. Each cell is assigned a state, higher score indicates better assignment of this cell to the respective state. **b,** UMAP of cells belonging to the geminal center (GC) maturation states including DZ and LZ. Color indicates the state. **c,** Ig light chain restriction. Color coded are either kappa or lamda Ig light chains.

## Materials and Methods

### Cell Culture

WTC-11 iPSCs (Coriell Institute for Medical Research – GM25256) were verified to display a normal karyotype, contamination free and regularly tested for mycoplasma. They were cultured in Essential 8™ Medium (E8) (Thermo Fisher Scientific, Cat. #A1517001) on Vitronectin XF™ (Stem Cell Technologies, Cat. #07180) coated tissue culture plates. Cells were maintained at 37 ° C at 5 % CO2. iPSCs were split using Accutase (StemCell Technologies - #07922) and E8 supplemented with 10 µM Y-27632 dihydrochloride (RI) (Tocris, Cat. No 1254). After dissociation into single cells, cells 1 volume of E8+RI was added, cells spun at 200 g for 5 min and resuspended and plated in E8+RI.

### SDR-seq in human iPSCs

WTC-11 iPSCs were dissociated into single cells using Accutase, filtered through a 40 µm cell strainer and counted. 1.5 x10^6 cells were transferred to a new 15 ml conical tube and spun at 500 g for 3 min.

For the glyoxal fixation condition the supernatant was removed, cells were resuspended in 200 µl of glyoxal solution fixation solution (3 % glyoxal, 20 % EtOH, 0.75 % Acetic Acid – Glacial, pH = 4.0) and incubated for 7 min at room temperature. 1 ml ice-cold wash buffer 1 (1x PBS with 2 % BSA, 1 mM DTT and 0.5 U/µl RNasin® Plus Ribonuclease Inhibitor - Promega #N2615) was added and cells were spun at 500 g for 3 min at 4 C. Supernatant was carefully removed and wash step was repeated with wash buffer 1 for a total of 2 washes. Cells were resuspended in 175 µl ice-cold permeabilization buffer (10 mM TRIS-HCl pH 7.5, 10 mM NaCl, 3 mM MgCl2, 0.1 % Tween-20, 0.2 U/µl RNasin® Plus Ribonuclease Inhibitor, 1mM DTT, 2 % BSA, 0.1 % IGEPAL CA-630 and 0.01 % Digitonin) and incubated for 4 min on ice. 1 ml of ice-cold wash buffer 2 (10 mM TRIS pH 7.5, 10 mM NaCl, 3 mM MgCl2, 0.1 % Tween-20, 0.2 U/µl RNasin® Plus Ribonuclease Inhibitor, 1mM DTT and 2 % BSA) was added and tube was gently inverted 4-6 times. Cells were spun at 500 g for 5 min at 4 C and resuspended in ice-cold resuspension buffer (1x PBS, 2 % BSA, 1 mM DTT and 0.2 U/µl RNasin® Plus Ribonuclease Inhibitor), filtered through a 40 µm strainer, counted and diluted to 1.4 x10^6 cells/ml.

PFA fixation conditions were followed as described elsewhere with adaptions (Rosenberg et al., 2018). In short, the supernatant was removed, cells were resuspended in 1 ml 1x PBS with 0.2 U/µl RNasin® Plus Ribonuclease Inhibitor and 3 ml of 1.3 % PFA solution (in 1x PBS) were added. Cells were fixed for 10 min on ice. 160 µl of permeabilization buffer (5 % Trition-X 100 with 0.2 U/µl RNasin® Plus Ribonuclease Inhibitor) was added, the tube gently inverted for 4-6 times and cells were incubated for 3 min on ice. Cells were spun at 500 g for 3 min at 4 C, supernatant was carefully removed and cells were resuspended in 500 µl of 1x PBS with 0.2 U/µl RNasin® Plus Ribonuclease Inhibitor. 500 µl of ice-cold 100 mM TRIS-HCl at pH 8.0 was added and mixed by inverting the tube. 20 µl of permeabilization buffer was added and mixed by inverting the tube 4-6 times. Cells were spun at 500 g for 3 min at 4 C, supernatant removed, resuspended in 300 µl of 0.5x PBS with 0.2 U/µl RNasin® Plus Ribonuclease Inhibitor, filtered through a 40 µm strainer, counted and diluted to 1.4 x10^6 cells/ml.

RT master mix consisting of a final concentration of 1x RT Buffer, 0.25 U/µl Enzymatics RNAse Inhibitor (Biozym – 180520), 0.2 U/µl RNasin® Plus Ribonuclease Inhibitor, 500 mM dNTPs and 20 U/µl Maxima H Minus Reverse Transcriptase (ThermoFisher - EP0752) was prepared on ice for in 8 µl for a total reaction volume of 20 µl. 4µl of RT oligo (12.5 µM) were combined in each 96-well plate with 8 µl RT master mix (Supplementary Table 1 and 2). 8 µl fixed and permeabilized cells (10000 cells total) were added to each well, yielding a total reaction volume of 20 µl. RT was performed in a thermocycler using the program below. All RT reactions were pooled into a 15 ml conical tube, 1 volume of ice-cold 1x PBS with 1 % BSA was added and cells were spun at 500 g for 5 min and supernatant was removed.

| Tempearture<br>°C | Time | Cycle |
| --- | --- | --- |
| 50 | 10 min |  |
| 8 | 12 sec | 3x |
| 15 | 45 sec |  |
| 20 | 45 sec |  |
| 30 | 30 sec |  |
| 42 | 2 min |  |
| 50 | 3 min |  |
| 50 | 5 min |  |
| 4 | - |  |

Samples were processed using the Tapestri microfluidic device from Mission Bio (MB51-0007, MB51-0010, MB51-0009) according to manufactures protocol with modifications. *In-situ* RT processed cell pellet from previous step was resuspended in the cell buffer of Mission Bio, cells were counted and diluted to the appropriate concentration of 3000-4000 cells/µl. Custom primers were used in the multiplexed droplet PCR amplification step. RNA primers were designed using the TAP-seq primer prediction tool with at targeted Tm of 60 ° C^4^. gDNA primers were designed using the Tapestri Designer (https://designer.missionbio.com). Version 1 gDNA and RNA primers both had CS and R2N overhangs (only used in POP experiment). Version 2 gDNA primers had CS and R2N, whereas RNA primers had CS and R2 overhangs. Forward and reverse stock primers had a concentration of 16 µM and 90 µM for both versions, respectively. For Version 1 final sequencing libraries were generated according to the Mission Bio user guide. For Version 2 RNA and gDNA sequencing libraries were generated separately using the corresponding library amplification primers (gDNA: R1N – R2N, RNA: R1N – R2).

### SDR-seq in primary non-Hodgkin lymphoma patient samples

The study (S-254/2016) was approved by University of Heidelberg’s Ethics Committee. Informed consent from every patient was gathered beforehand. Lymph node samples were processed and frozen following previously described methods^45,46^. Frozen samples were thawed, added to 10 ml RPMI (Gibco, #11875093) supplemented with 10 % FBS and 0.5 mM EDTA and spun at 400 g for 5 min. Cells were resuspended in 10 ml 1x DPBS supplemented with 5 % FBS, filtered through a 70 µm strainer and spun at 400 g for 5 min. Cells were resuspended in 100 µl bead solution from the Dead Cell Removal Kit (Miltenyi Biotec, #130-090-101) and incubated for 15 min in the dark. Binding buffer was prepared according to manufactures protocol, and the LS column (Miltenyi Biotec, #130-042-401) washed with 500 µl binding buffer. Cells were applied to the column and washed 4x with binding buffer while collecting the flow through. Cells were spun at 400 g for 5 min and resuspend in 1 ml of binding buffer. We proceeded with glyoxal fixation and SDR-seq as described above.

### SDR-seq data analysis

For each SDR-seq dataset generated, we first performed custom barcode identification and error correction, mapping of reads to custom reference sequences, and the building of read and deduplicated UMI matrices. This was performed with a software package we named SDRranger (https://github.com/hawkjo/SDRranger).

The full barcode structure for the RNA targeted libraries is of the format cell barcode 1 – Variable length linker (14-17 bp) – cell barcode 2 – Constant length linker (15 bp) – Sample BC – UMI. The gDNA libraries are the same but lacking the final sample BC and UMI. To identify these, we first aligned each read to all possible linker backbone sequences to account for the variable length linker sequences. We discarded alignments with length-normalized alignment scores more than two standard deviations below average, measured from the first 10,000 reads. We then performed error correction on BC1 and BC2 to unique corrected barcodes with Levenshtein distance zero or one. Due to the adjacent UMI, the Sample BC does not have an identifiable end point in the case of insertions and deletions, so we adapted the method in J. A. Hawkins et al., to correct to Sample BCs with Free Divergence zero or one, and with no other BC with Free Divergence only one higher^47^.

Following barcode identification, reads are mapped to custom alignment references built for each gDNA and RNA library. For gDNA the chromosomal locations of the amplicons were used to extract reference sequences. For RNA the site of the primer binding until the end of the polyA was used to extract reference sequences. References sequences were extracted from the GRCh38 genome assembly. Custom Fasta and gtf files were generated and used to build references using the genomeGenerate function of STAR (v2.7.11a). Separate gDNA or RNA sequencing libraries were aligned to the corresponding reference, except for checking the specificity of the sequencing libraries (Extended Data Fig. 3b).

For the POP experiment, reads were separated into gDNA and RNA reads before barcode identification by a separate mapping step to the corresponding references. Final BAM files are produced which contain tags with cell barcode and sample barcode sequences for each read, both before and after error correction.

Matrices were then constructed by tallying reads by cell barcode and sample BC vs gene or gDNA amplicon. To construct the UMI matrices, UMIs were deduplicated by adapting the directional network deduplication method described in the UMItools package^48^. For all reads from a given cell and given gene or amplicon, a connectivity graph of all observed UMIs is constructed. Each node is a unique UMI sequence and read count of that sequence, and directed edges are added between nodes A and B if the two UMIs have a Free Divergence of one, to allow for indels, but only if n_A_ ≥ 2n_B_ – 1 reads, where n_A_ and n_B_ are the respective number of reads. This is based on the observation that each additional error to a UMI sequence should reduce the frequency of observing that sequence. Furthermore, only one incoming “parent” edge is allowed per node, to avoid artifactual connections through singletons. The final number of UMIs is the number of connected components of the graph at the end of this process. This is repeated for each cell and sample barcode for each gene or amplicon to build the full matrix.

The resulting cell-gene (UMIs - RNA) and cell-amplicon (reads - gDNA) matrices were analyzed in R using the Seurat R package (v5.0.3). First a general threshold per cell was set on reads/cell (RNA+gDNA) based on rank-rank plots of reads/cell to ranked cells to determine an initial set of cells to include in the analysis. Then multiple metrics including detected genes, detected gDNA amplicons, number of UMIs (RNA), number of reads (gDNA) and the ratio of correctly mapped sample BC reads per cell were used to filter for high-quality cells. Contaminating reads that did not belong to the maximum sample barcode found per cell were removed. RNA count matrixes were processed using the Seurat R package (log-normalize per, scaled). Principial component analysis (PCA) was performed on all genes measured with subsequent UMAP embedding. For clustering the shared nearest neighbor graph was calculated and used as input for Louvain clustering.

Each cell that was defined as high-quality was then used to call variants using the GATK HaplotypeCaller (v4.2.3.0)^49^. Individual bam files were generated using the cell barcode of the high-quality cells using the package sinto (v0.10.0). Each induvial cell bam was modified to contain the cell barcode in the read name, was indexed using samtools (v1.17) and the MAPQ scores were set from 255 (STAR output) to 60 and to be compatible as an input in the GATK HaplotypeCaller. GATK Haplotype caller was run using no maximum read threshold per cell and using a diploidy of two, resulting vcf files were merged to yield a matrix of cells to variants compatible as input for Seurat. Low frequency variants (< 0.1 % for editing, < 0.3 % for primary patient samples) were removed, remaining variants were input into the Ensembl Variant Effect Predictor for functional annotation^50^. This functional annotation was added as metadata to the Seurat object. Genotypes of the GATK Haplotype caller were added as an assay to the previous Seurat object, while the remaining output was added as metadata. WT alleles were included based on the read depth for a given amplicon per cell. Both WT alleles and variant alleles were excluded from subsequent analysis if read depth was low (< 5 reads) or the genotype quality score of the GATK HaplotypeCaller was low (< 30 GQ).

### Cloning, molecular biology and generating transgenic iPSCs

For the constitutive CRISPRi, PEmax and PEmax-MLHdn1 cell lines the corresponding transgene was inserted into the AAVS1 locus in WTC-11 iPSCs as previously described using specific TALENs^5,51^. The AAVS1 targeting vector containing the homology arms, the CAGG promotor and a WPRE was a kind gift from J. A. Knoblich (Institute of Molecular Biotechnology of the Austrian Academy of Science -IMBA, Vienna BioCenter, Vienna, Austria). For the CRISPRi plasmid the pHR-UCOE-SFFV-dCas9-mCherry-ZIM3-KRAB (Addgene, #154473) was modified to pHR-UCOE-SFFV-dCas9-mCherry-KRAB-MECP2 with DNA fragment ordered from Twist Bioscience containing KRAB-MECP2. dCas9, KRAB-MECP2 and dTomato were amplified and cloned into the AAVS1 targeting vector described above with the NEBuilder® HiFi DNA Assembly Master Mix (NEB, #M5520). For the PEmax and PEmax-MLHdn1 plasmid the CRISPRi plasmid was used as a backbone while inserting the PEmax or PEmax-P2A-MLHdn1 (Addgene, #174828) sequences with the NEBuilder® HiFi DNA Assembly Master Mix. To generate transgenic iPSCs expressing the CRISPRi, PEmax or PEmax-MLHdn1 transgene from the AAVS1 locus, WTC-11 iPSCs were electroporated with the corresponding homology plasmid (3 µg per electroporation) and two TALEN plasmids (0.75 µg per electroporation each) targeting the AAVS1 locus (Addgene, #52341 and #52342). iPSCs were dissociated into a single cell suspension, counted and 1×10^6^ cells electroporated using the CB-150 program of the 4D-Nucleofector™ System and the P3 Primary Cell 4D-Nucleofector® X Kit L (Lonza, #V4XP-3024) according the manufactures protocol and plated in E8+RI. Cells were sorted 7-10 days after the electroporation for dTomato, plated at low density in E8+RI, grown to colonies, picked and genotyped. Positively genotyped clones were checked for homogenous dTomato signal, validated for activity in a corresponding assay and three clones of each cell line were subjected to a genotyping array screening using a Infinium Global Screening Array-24 Kit (Illumina, #20030770) to check for chromosomal rearrangements in iPSCs clones. Only clones that showed no or minor differences to the WTC-11 WT parental cell line were used in this study.

### gRNA/pegRNA design and library cloning

All gRNA and pegRNA libraries were cloned in pools. eQTLs were selected based on high confidence from published data and both lowly expressed genes (CPM > 150) and essential genes in iPSCs removed also based on published data^5,52–54^. eQTLs for the base editor screen were further filtered by overlap with ATAC-seq peaks and compatibility with transversion by adenine or cytosine base editors (A>G, C>T, G>A, T>C)^55^. Sites to introduce STOP codons were chosen manually in the selected genes. pegRNAs were designed using PrimeDesign (https://drugthatgene.pinellolab.partners.org) and linkers to separate the pegRNA from the tevopreQ1 3’ stabilizing sequence were designed using pegLIT (https://peglit.liugroup.us)^56,57^. For the CRISPRi experiment the spacer sequences of the above pegRNAs were used for eQTLs and STOP controls, while gRNAs targeting the TSS for genes predicted to be affected by eQTLs were designed using CRIPSRick (https://portals.broadinstitute.org/gppx/crispick/public) and NTCs were chosen from the GeCKO-v2 library^58–60^. Base editing gRNAs were designed and selected based on highest predicted editing efficiency using the BE-Hive tool^61^. The pegRNA screening vector was a kind gift from J. A. Knoblich (Institute of Molecular Biotechnology of the Austrian Academy of Science -IMBA, Vienna BioCenter, Vienna, Austria). It was modified to remove the ERT2-Cre-ERT2 sequence and the gRNA scaffold and include a 3’ stabilizing tevopreQ1 after the insertion site for pegRNAs via BbsI Golden Gate cloning. The base editing vectors were all-in-one cytosine or adenine base editor + guide expression constructs (Addgene, #158581 and #179097). The gRNA screening vector was a modified CROP-seq vector (Addgene, #86708) to also express an EGFP and include a capture sequence in the scaffold of the gRNA^62^. Oligos for pegRNA and gRNA libraries were order from IDT as oPools. pegRNA oligos included a spacer sequence, PBS and RT with overhangs for amplification that included BbsI sequences compatible with Golden Gate cloning. Spacer and PBS/RT sequences were separated by a constant sequence containing two Esp3I sites for a second round of Golden Gate cloning to introduce the pegRNA scaffold. gRNA oligos consisted of spacer sequences and overhangs for amplification that included BbsI sequences compatible with Golden Gate cloning. Oligos were amplified (8 cycles) with compatible primers. The purified PCR product was cloned into the respective pegRNA or gRNA screening vector described above using BbsI and Golden Gate cloning. Electrocompetent bacteria (Lucigen, #60242-1) were electroporated (10 µF, 600 Ohms, 1800 V, E = 184 V/cm) with purified ligation product and grown in a pool for 10 hours at 30 ° C before extracting plasmid DNA. For pegRNA and base editor guide libraries a scaffold sequence was ordered with overhangs that included Esp3I overhangs (IDT), amplified with complementary primers (8 cycles), purified and cloned as described above using Esp3I Golden Gate cloning.

### Virus production, infection of human iPSCs and lipofection of human iPSCs

Lentiviruses were produced in HEK293 grown in DMEM supplemented with 10 % FBS, 1x GlutaMAX™ (Gibco, #35050061), 100 U/ml Penicillin-Streptomycin (Gibco, # 15140122) and 1x MEM Non-Essential Amino Acids (Gibco, # 11140050) and coated using VSV-G. Day before transfection HEK293 cells were plated at 80 % confluency, plasmids were lipofected using Lipofectamine™ 3000 Transfection Reagent (L3000001) and split 1:10 five hours after lipofection. Supernatant was collected three days after lipofection, cell debris pelleted at 200 g for 5 min at 4 ° C, remaining supernatant spun at 28,000 g for five hours and virus pellet resuspended in the appropriate volume of E8+RI. On the day of infection, human iPSCs were split 1:2.5 two hours before the infection using Accutase. Infections were performed overnight in E8+RI. Media was replaced the next day with E8. For some editing experiments constructs were only expressed transiently in human iPSCs. For this transfection was done using the Lipofectamine Stem Transfection Reagent (STEM00003) according to manufactures protocol.

### Target selection and subsampling for panel size testing experiment

Candidate cis-regulatory elements (cCRE) for five human iPSC lines (H1, H7, H9, iPS DF 6.9, iPS DF 19.11) were obtained from SCREEN (https://screen.encodeproject.org) and corresponding regulatory elements subset from these^26^. Genomic regions were defined as OEG or NOEG if the gene overlapping that genomic region was expressed in bulk RNA-seq data (> 10 CPM)^5^. 120 regions were randomly subsampled for each cCRE within the OEG and NOEG classes and primers designed as described above. Each cCRE with OEG and NOEG was equally represented in both the total and shared panels. Genes were subset into highly (> 400 CPM), medium (< 400, > 40) and lowly (< 40, > 4) expressed genes. Primers were designed as described above. After determining high-quality cells in all panels as described above, they were subset from the bam file and reads/cell for gDNA and RNA was scaled according to panel size in a way so the average number of reads/cells for shared gDNA and RNA targets was the same.

### Maturation state assignment in primary tumor samples and Ig light chain restriction analysis

B-cell maturation states were mapped to each tumor sample from a published reactive lymph node single-cell RNA sequencing dataset through shared gene expression features using previously described methods^33,35^. Gene expression was used to determine Ig light chain restriction. Log-normalized counts (without batch-effect correction to prevent bias introduced by sample integration) were used to find transfer anchors and project samples on the reference PCA (50 dimensions) and UMAP (2 dimensions) reductions. Expression of immunoglobulin kappa (IGKC) and lambda (IGLC1-7) light chain genes was used to determine cells’ malignancy through light chain restriction^36^.

### Data analysis for primary non-Hodgkin lymphoma patient samples

Separate NGS sequencing libraries for gDNA and RNA were analyzed with SDRranger and variants were called as described above. Low frequency variants (< 5 % or less than 30 heterozygous/homozygous variants) were excluded from the analysis. GO term analysis was performed using the R package topGO (v2.54.0) using the weigth01 algorithm and Fisher’s exact test. Enrichment for BP was computed for the top 21 DE genes in LZ vs DZ across all patient samples versus all genes measured.

### Statistics and reproducibility

No data was excluded from analysis, cutoffs for defining high-quality cells in SDR-seq were set as described above. Differential gene expression testing in the single cell data was performed using MAST and by sub-setting cells in the respective genotype within given cell or perturbation states^63^. Differential abundance testing of variants between maturation states in primary non-Hodgkin lymphoma patient samples was performed using chi-squared testing followed by adjusting *P*-values by the Benjamini–Hochberg method.

## Data Availability

Raw sequencing data and processed data will be uploaded on GEO and EGA. Accession codes will be made available upon publication.

## Code Availability

All relevant code will be deposited on GitHub upon publication. Code containing SDRranger to generate count/read matrices from RNA or gDNA raw sequencing data is available under https://github.com/hawkjo/SDRranger.

